# Conserved ancillary residues situated proximally to the VIM-2 active site affect its metallo β-lactamase activity

**DOI:** 10.1101/2024.09.18.613613

**Authors:** Diamond Jain, Tejavath Ajith, Jyoti Verma, Debasmita Chatterjee, Anindya S. Ghosh

## Abstract

Verona-integron-metallo-β-lactamase (VIM-2) is one of the most widespread class B β-lactamase, responsible for β-lactam resistance. Although active-site residues help in metal binding, the residues nearing the active-site possess functional importance. Here, to decipher the role of such residues in the activity and stability of VIM-2, the residues E146, D182, N210, S207, and D213 were selected through in-silico analyses and substituted with alanine using site-directed mutagenesis. The effects of substitution mutations were assessed by comparing the changes in the β-lactam susceptibility pattern of E. coli host cells expressing VIM-2 and its mutated proteins. VIM-2_N210A enhanced the susceptibility of the host by ∼4-8 folds against penicillins and cephalosporins while the expression of VIM-2_D182A radically increased the susceptibility of the host. However, expression of VIM-2_E146A reduced the susceptibility of the host by 2-fold. Further, proteins were purified to homogeneity, and VIM_N210A and VIM_D182A displayed reduced thermal stability than VIM-2. Moreover, in vitro catalytic efficiencies of VIM-2_D182A were drastically reduced against all the β-lactams tested whereas the same were moderately reduced for VIM-2_N210A. Conversely, the catalytic efficiency was marginally altered for VIM_E146A. Overall, we infer that both N210A and D182A substitutions negatively affect the performance of VIM-2 by influencing substrate specificity and stability, respectively.

## Introduction

Metallo β-lactamases (MBLs) are one of the most prominent enzymes responsible for the inactivation of a wide range of β-lactam antibiotics that leads to antimicrobial resistance, which is a matter of concern for antimicrobial therapy. Unavailability of clinically useful inhibitors against these MBLs is limiting the therapeutic options (Reddy, *et al*. 2020). MBLs are included in class B of Ambler classification system, which is further divided into three subclasses (B1, B2, and B3) based on their primary sequences and different active-site features (Hall and Barlow 2005). Most of them belong to subclass B1, among which imipenemase (IMP), Verona integron metallo β-lactamase (VIM), and New Delhi-metallo β-lactamase (NDM) families are widely disseminated (Yang, *et al*. 2023). Among all the three enzymes, VIM type metallo β-lactamases are significantly prominent and there are various reports on the presence of *bla*_VIM-2_ in clinical and environmental isolates of different Gram-negative bacteria (Papa-Ezdra, *et al*. 2023, Yum, *et al*. 2002).

VIM-2 was first reported from an isolate of *Pseudomonas aeruginosa* in France and then in Greece (Mavroidi, *et al*. 2000, Poirel, *et al*. 2000). The coexistence of *bla*_VIM-2_ and *bla*_NDM-1_ in the clinical isolate of *P. aeruginosa* is reported from India (Paul, *et al*. 2016). It hydrolyses a broad range of the β-lactams, namely, penicillins, cephalosporins, cephamycins, oxacephamycins and carbapenems but not the monobactams (Poirel, *et al*. 2000). Structurally, VIM-2 (29.7 kDa) is a 266 amino acid long protein with about 32% similarity with BcII; 31% with IMP-1; 27% with CcrA; 24% with BlaB; 24% with IND-1; 21% with CphA-1; 32.4 % with NDM-1; and 11% with L-1 (Lee, *et al*. 2009, Poirel, *et al*. 2000). It differs from VIM-1 by seventeen amino acid residues. Similar to other sub-class B1 MBLs, VIM-2 possesses two zinc ions sandwiched within the conserved αβ/βα conserved motif of metal hydrolases for the catalysis. The Zn1 is coordinated with H114, H116, H179 and Zn2 is coordinated with D118, C198 and H240 (Meini, *et al*. 2014).

Apart from the active-site, the residues lying near or far away from it play an imperative role in the activity and stability of metallo β-lactamases (Bahr, *et al*. 2021). Moreover, some of these conserved residues may impart altered functions in different MBLs. Therefore, in this work we aimed to identify the residues that are lying near or far away from the active-site which can affect the activity and the stability of VIM-2. The five residues, *viz.*, E146, D182, S207, N210 and D213 conserved in NDMs and VIM-2 were selected and substituted with alanine to nullify the charge without affecting the secondary structures. The effect of these substitutions was estimated by assessing the change in susceptibility pattern of the expression host against β-lactam. Further, the wild type and the mutated proteins were purified, and assayed for their *in vitro* activity in terms of catalytic efficiency, thermal stability, and effect on the secondary structures.

## Materials and Methods

### Bacterial strains, plasmids, media and growth conditions

*P. aeruginosa* isolate harboring *bla*_VIM-2_ was received as a gift (Paul, *et al*. 2016). The full length *bla*_VIM-2_ was cloned in the tightly regulated arabinose inducible pBAD18cm (Guzman, *et al*. 1995) vector and the truncated gene devoid of signal peptide was cloned in T7 promoter based pET28a vector (Novagen, Madison WI, USA). The *in vivo* studies were carried out in *E. coli* CS109 (Ghosh and Young 2003) and expression for purification was done in *E. coli* BL-21(DE3) (Stratagene, West Cedar Creek, TX, USA). Both plasmids were maintained in *E. coli* XL1-Blue (Stratagene, La Jolla, CA, USA). Bacterial strains were grown and maintained in Luria Bertani broth and agar at 37°C for 16-18 h. Minimum inhibitory concentrations of antibiotics were determined in cation adjusted Mueller Hinton broth (Hi-media, Mumbai, India). The primers used in this study are mentioned in table S1. The reagents for molecular biology works were procured from New England Biolabs (Ipswich, MA, USA). Unless otherwise specified, all the other reagents and antibiotics were purchased from Sigma-Aldrich (St. Louis, MO USA).

### Cloning and site-directed mutagenesis

The full-length gene *bla*_VIM-2_ was amplified using the genomic DNA of clinical isolate as template and cloned in pBAD18cm vector under *Nhe*I and *Hin*dIII restriction sites. The β-lactamases are localized in the periplasm of bacteria. Therefore, the signal peptide from N-terminal of VIM-2 was predicted using SignalP4.0 (Petersen, *et al*. 2011) and the primers were designed by omitting the nucleotides coding for the signal sequence. The truncated form of *bla*_VIM-2_ (devoid of signal peptide) was cloned in pET28a vector under *Nde*I and *Hin*dIII restriction sites for soluble cytoplasmic expression and purification. Site-directed mutagenesis was carried out using QuikChange site-directed mutagenesis kit as per the manufacturer instructions (Agilent Technologies, CA, US) and all the mutations were confirmed by sequencing (Eurofins Pvt. Ltd., Bangalore, India).

### Expression of gene, protein production and purification

To study the *in vitro* activity, the proteins were purified using 6X-His tag Ni^2+^-affinity chromatography, since the presence of 6X-His tag have negligible effect on the activity of VIM-2 (Makena, *et al*. 2016). *E. coli* BL-21(DE3) cells were transformed with the pET28a recombinant constructs, and optimized for the gene expression and protein production using 0.5 mM IPTG as an inducer of gene expression at 16 °C for 16 h. The bacterial cultures harbouring the proteins generated were harvested by centrifugation at 10000 g for 10 min, resuspended in lysis buffer and lysed by using sonication. The supernatants containing the expressed proteins were separated by centrifugation at 50000 g for 90 min, and loaded on the 1 ml HisTrap™ column and purified using AKTA prime plus system as previously described (Kumar, *et al*. 2020). The purified proteins (Fig. S1) were analysed through 12 % SDS-PAGE and the most homogenous protein elutes were pooled together. Imidazole was removed by dialysis using three subsequent changes in buffer (10mM Tris, 150 mM NaCl, pH 7.8) and concentration of the proteins was estimated using Bradford assay.

### Antibiotic susceptibility assay

Antibiotic susceptibility of *E. coli* CS109 harbouring VIM-2 and its mutants were determined to assess the effect of substitutions on the *in vivo* activity of VIM-2. For this, *E. coli* CS109 cells were transformed with recombinant pBAD18cm constructs of VIM-2 and its mutants. The minimum inhibitory concentrations of β-lactam antibiotics were determined through micro-broth dilution method using 96 well plates. Antibiotics were serially diluted by two-fold with 30 µL of MH Broth. Concentrations of antibiotics were in the range between 2048 to 2.0 μg mL^-1^ for penicillins, 256 to 0.5 μg mL^-1^ for cephalosporins and 32 to 0.125 μg mL^-1^ for carbapenems. Total volume of MH broth in each well was made up to 300 μL by adding overnight grown culture at a concentration of C 10^5^ cells per well and incubated at 37°C for 16-18 h. The optical density of culture was measured at 600 nm and results were interpreted as per CLSI guidelines (Wayne 2014). The experiments were conducted thrice with three replicates for each set of experiment.

### *In vitro* enzyme activity analysis and enzyme kinetics

The activity of the purified proteins was assessed using nitrocefin hydrolysis assay. The hydrolysis of nitrocefin (100 µM) by purified proteins (100 nM) in buffer (10mM Tris-HCl, 150 mM NaCl, pH 7.8) was monitored spectroscopically at *A*_490nm_. Zinc sulphate was supplemented at a final concentration of 1µM. The enzyme kinetics parameters, *K_m_*, *k*_cat_ and *k*_cat_/*K_m_*were calculated to determine the *in vitro* catalytic efficiency of the purified proteins using steady state kinetics. The concentrations and different parameters for the substrates and enzymes used in this study are mentioned in the table S2. The change in substrate concentration after hydrolysis was measured spectrophotometrically at 25 °C in 10 mM Tris-HCl buffer (pH - 7.8) supplemented with 1 µg/mL BSA and 1 µM ZnSO_4_ at an interval of 5 sec for about 1-2 min. The initial rate of reaction at different substrate concentrations were calculated, and were plotted against substrate concentrations of the antibiotic and analyzed using online curve-fitting tool (Kumar, *et al*. 2020). The *k_cat_* values were obtained by dividing *V*_max_ by total [E].The values reported are the mean of three independent replicates.

### Determination of thermal stability using differential scanning fluorimetry

The thermal stability of the purified proteins was estimated by determining the melting temperatures using differential scanning fluorimetry (DSF). The reaction was set up using 5 µg proteins in 10 mM Tris-HCl pH 7.8 containing 150 mM NaCl and 50 µM Zinc sulphate. The 5X final concentration of SYPRO Orange dye was added to each well. Finally, the fluorescence readings (monitored at 492 nm excitation and 610 nm emission) were measured using BioRad CFX-96 qPCR system (BioRad, Hercules, California) between 20°C and 90°C, by increasing the temperature linearly in steps of 0.5°C every minute. The experiments were performed thrice in triplicates and the difference in the melting temperatures obtained were ±0.5°C.

### Secondary structure assessment using circular dichroism

Circular dichroism spectroscopy is usually employed to check the secondary structural integrity of the native and mutated proteins. The far UV (190-240nm) circular dichroism (Far UV CD) spectrum of the proteins was determined using a Jasco J-1500 spectropolarimeter (JASCO International Pvt. Ltd., Japan). The spectral data was collected by placing the sample in a quartz cell with a path length of 0.1 cm, at 25 °C with a 0.1 nm step resolution, 1 s time constant, 10 milli-degree sensitivity at 1.0 nm spectral band-width, with a scanning speed of 50 nm min^-1^. To reduce the noise and random instrumental error, an average spectrum of four successive accumulations were taken over 190–260 nm range and the corrected spectra were obtained by subtracting the baseline buffer spectrum. All the proteins samples were diluted accordingly for the CD spectroscopy. The obtained ellipticity values (θobs; in degrees) were converted to mean residue ellipticity (MRE) in degrees per square centimeter per decimole (mdeg cm^2^ dmol^-1^) using the following equation:

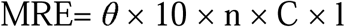

where θ is the observed ellipticity (in degrees), n is the total number of amino acid residues in the protein, C is the molar concentration of protein and l is the path length (in centimeters). Further, the MRE values at 222 nm were used to calculate the helical content using the following relations as described by (Chen, *et al*. 1972) and adopted in (Khan and Rehman 2016).

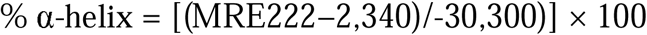

## Results

### Insights into the structural features of VIM-2 for selection of mutation sites

Metallo β-lactamases are widely studied for understanding the role of non-active-site residues in maintaining the catalytic activity and the thermal stability of these robust enzymes (Bahr, *et al*. 2021). These residues can be numbered according to the primary sequence as well as class B β-lactamase (BBL) numbering, based upon the three-dimensional structural alignment of the MBLs (Garau, *et al*. 2004). Previously, the importance of the residue E152 (BBL E149) was discussed for its role in the activity of NDM-5 and NDM-7 metallo β-lactamases (Kumar, *et al*. 2020, Kumar, *et al*. 2017a). NDM-9 is a natural variant of NDM-1 with E152K (BBL E149K) substitution, which imparts higher hydrolysing activity towards carbapenems and cephalosporins (Wang, *et al*. 2014). This residue is conserved in VIM-2 at position 146 (BBL 149) (Fig. 1). Moreover, two distinct VIM variants, VIM-7 and VIM-32 also possess E146A substitution along with the other natural substitutions (Samuelsen, *et al*. 2008). The residue S207 (BBL S230) in VIM-2 is similar to S217 of NDMs and lies nearer to the substrate binding site. The S217A substitution in NDM-1 and NDM-4 has led reduction in the activity of the enzyme (Ali, *et al*. 2021, Verma, *et al*. 2023). The conserved residue N210 (BBL N233) interacts with the zinc ion and proposed to form an oxyanion hole along with zinc for accommodation of the carbonyl oxygen of the hydrolysed β-lactam ring (Docquier, *et al*. 2003). The residue D213 (BBL 236) also lies closer to the active-site and zinc interacting residues. Accordingly, the residues E146, D182, S207, N210 and D213 (as shown in Fig. 1) were substituted with alanine, individually and assayed for the impact of these substitutions on the susceptibility pattern of VIM-2. The mutants exhibiting reduced activity were further purified to analyse their effect on *in vitro* catalytic activity and thermal stability.

**Fig. 1.**
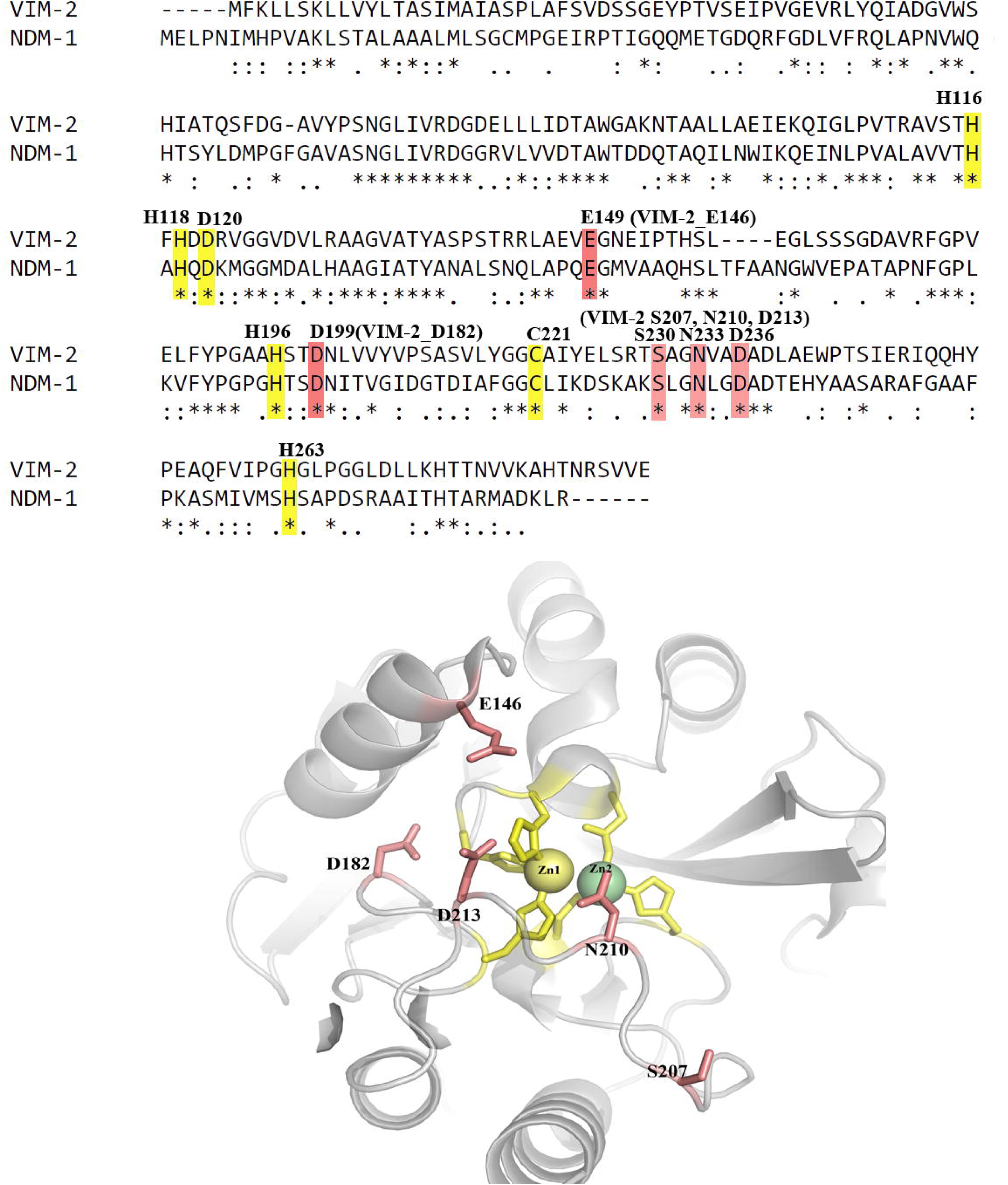
Insight into the structure of VIM-2 for the selection of amino acid residues for mutagenesis. The yellow colour depicts the active site residues bound to Zn1 and Zn2 of the protein, while the salmon red colour highlights the residues selected for mutagenesis. (PDB ID 4BZ3)

### E146A, D182A and N210A substitution altered the β-lactam susceptibility pattern of *E. coli* host expressing VIM-2

The susceptibility of *E. coli* CS109 cells harboring VIM-2 and its mutants towards different β-lactam antibiotics was assessed by determining MICs. To accomplish this, CS109 cells were transformed with the full-length constructs cloned in pBAD18cm and the expression of protein in the bacterial culture was induced with 0.02% arabinose. Ectopic expression of VIM-2 substantially (at least 64-fold) reduced the susceptibility of *E. coli* cells (Table 1) against all the tested antibiotics. Alanine replacement at the positions D182 and N210 dramatically reduced the activity of VIM-2 in *E. coli*. Ectopically expressed VIM-2_D182A could not resist activity of all the tested β-lactams, as shown by the susceptibility pattern that is nearly equal to that of *E. coli*. Cells expressing VIM-2_N210A displayed >4-fold increased sensitivity towards penicillins and cephalosporins as compared to VIM-2. The substitution at the position N210 with alanine had the most prominent effect on the susceptibility of cefotaxime where the cells carrying the mutated proteins showed a reduction of resistance by 64-fold as compared to the cells expressing wild-type VIM-2. However, no change in the susceptibility pattern was observed towards the carbapenems for VIM-2_N210A. Interestingly, cells expressing VIM-2_E146A exhibited decreased susceptibility by 2-fold towards all the tested β-lactams than that of the cells expressing compared to VIM-2. Therefore, VIM-2_D182A, VIM-2_N210A and VIM-2_E146A were further purified to understand their effects on *in vitro* activity of VIM-2. On the other hand, susceptibility pattern of the cells expressing S207A and D213A exhibited negligible alterations and accordingly, these residues were not considered for the further study.

**Table 1.** Minimum inhibitory concentrations of tested beta-lactam antibiotics against *E. coli* strain expressing VIM-2 and its mutants.

| Antibiotics | CS109 | MIC (µg/mL) |  |  |  |  |  |
| --- | --- | --- | --- | --- | --- | --- | --- |
|  |  | VIM-2 and its mutated variants |  |  |  |  |  |
|  |  | VIM-2 | E146A | D182A | S207A | N210A | D213A |
| *Penicillins* |  |  |  |  |  |  |  |
| Ampicillin | 16 | 1024 | 2048 | 16 | 1024 | 256 | 1024 |
| Ampicillin+EDTA* | 16 | 16 | 16 | 16 | 16 | 16 | 16 |
| Ticarcillin | 8 | 1024 | 2048 | 8 | 1024 | 124 | 1024 |
| Amoxicillin | 8 | 1024 | 2048 | 8 | 1024 | 256 | 1024 |
| *Cephalosporins* |  |  |  |  |  |  |  |
| Cephalothin | 4 | >512 | 1024 | 4 | >512 | 256 | 512 |
| Cefadroxil | 8 | 512 | 1024 | 8 | 512 | 64 | 512 |
| Cefoperazone | 4 | 512 | 1024 | 4 | 512 | 64 | 512 |
| Cefotaxime | 2 | 256 | 1024 | 2 | 128 | 4 | 256 |
| Cefotaxime+EDTA* | 2 | 2 | 2 | 2 | 2 | 2 | 2 |
| *Carbapenems* |  |  |  |  |  |  |  |
| Imipenem | 0.25 | 32 | 64 | 0.5 | 32 | 32 | 32 |
| Imipenem+EDTA* | 0.25 | 0.25 | 0.25 | 0.25 | 0.25 | 0.25 | 0.25 |
| Meropenem | 0.5 | 32 | 64 | 1 | 32 | 32 | 32 |
| Doripenem | 0.5 | 32 | 64 | 1 | 32 | 32 | 32 |

\*EDTA was used at a concentration of 400µM

### *In vitro* catalytic efficiencies of VIM-2 and its mutants substantiate the antibiotic susceptibility pattern

To assess the catalytic efficiency and the thermal stability of VIM-2 and its mutants, the proteins were purified using 6X-his tag Ni^2+^-affinity chromatography (Fig. 2a). The presence of his-tag has minimal effect on the activity of VIM-2 as reported previously (Makena, *et al*. 2016). The availability of zinc in purification media and buffers and the substitution mutations might affect the zinc content of the purified proteins (Docquier, *et al*. 2003). Therefore, the effect of excess zinc supplementation (1 µM) on the activity of the proteins (100nM) was assayed by the nitrocefin hydrolysis assay (Fig. 2b). VIM-2 efficiently hydrolysed nitrocefin (100 µM) as displayed by the instant colour change after 1 min of incubation and addition of excess zinc enhanced its activity. The zinc deficient VIM-2_D182A displayed minimal colour change for the nitrocefin hydrolysis and the addition of zinc in excess enhanced its activity to some extent. The N210A substitution also reduced the ability of VIM-2 to hydrolyse nitrocefin while upon addition of excess zinc, the hydrolysing ability was increased nearly to the levels of VIM-2. Interestingly, E146A substituted VIM-2 exhibited reduced activity as compared to VIM-2. However, addition of zinc significantly increased its activity. Since we intended to compare the effects of mutations with the wild type protein, further, the catalytic efficiencies, thermal stability and secondary structures of the purified proteins against various β-lactam substrates were determined in presence of excess zinc.

**Fig. 2.**
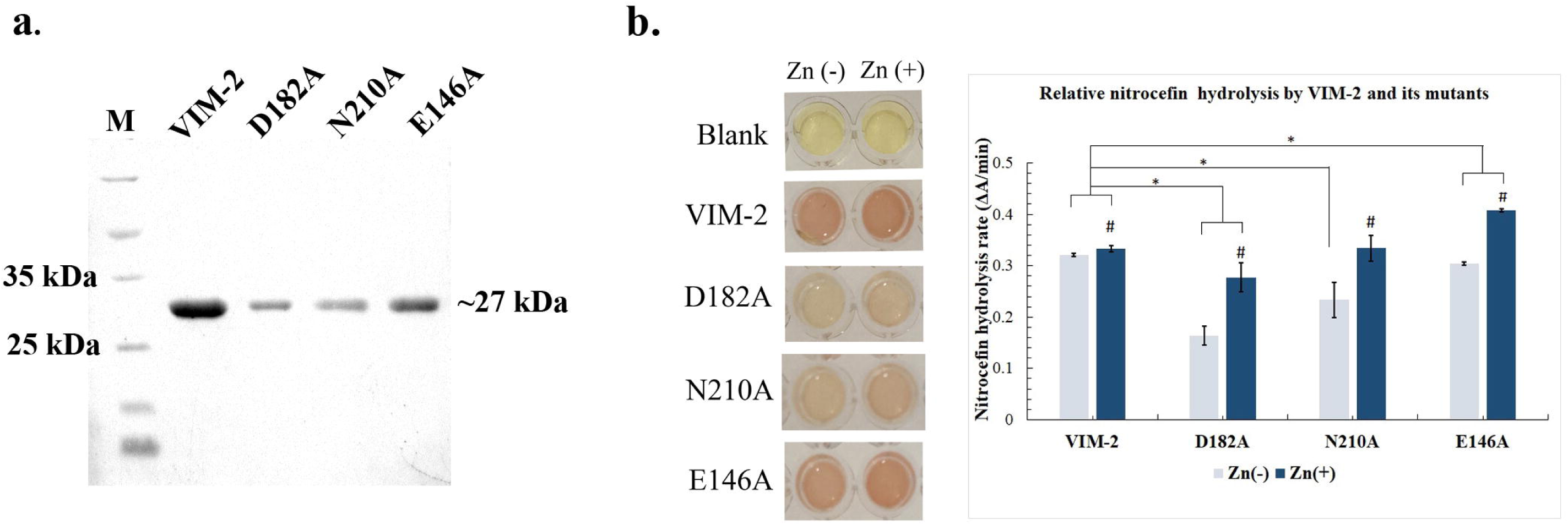
Purification and activity assessment of VIM-2 and its mutants using nitrocefin hydrolysis assay showing a. Purified homogenous eluted of VIM-2 and its mutants using Ni^2+^-affinity chromatography b. Colour change of nitrocefin from yellow to red after hydrolysis by VIM-2 and its mutants. Colour intensity can be directly corelated with the β - lactamase activity. Bar graph depiction of nitrocefin hydrolysis where ‘#’ represents significant change in activity of proteins at *p*<0.05 after supplementation with zinc, while ‘*’ significant change in activity of the mutants as compared to VIM-2 at *p*<0.05, determined using unpaired, two-tailed *t*-test.

To assess the *in vitro* catalytic efficiencies against various β-lactam substrates, the enzyme kinetics parameters (*k*_cat_, *K*_m_ and *k*_cat_/*K*_m_) of the purified proteins were determined (Table 2). Since, the supplementation of zinc enhanced the activity of the purified proteins; catalytic efficiencies of VIM-2 and all its mutants were determined in the presence of zinc (1 µM). VIM-2 effectively hydrolyzed all the β-lactam substrates tested. The catalytic efficiency of the D182A substituted VIM-2 was substantially reduced by >20-fold for the antibiotics belonging to penicillin group, namely, ampicillin and ticarcillin. Similarly, VIM-2_D182A exhibited about >10-fold reduction in catalytic efficiency for the antibiotics from cephalosporin group, *viz.*, cephalothin and cefotaxime whereas a >50-fold reduction was observed for the antibiotics belonging to the carbapenems, e.g., imipenem and meropenem. Further, N210A substituted VIM-2 displayed variations in the hydrolysis of the substrates and corroborated with the antibiotic susceptibility pattern. For penicillins, the N210A substituted VIM-2 was able to hydrolyze ampicillin which was comparable to VIM-2, while about 2-fold reduction in the catalytic efficiency was observed for ticarcillin. VIM-2_N210A exhibited nearly 2-4-fold reduction in the catalytic efficiency for cephalothin and cefotaxime. As observed for antibiotic susceptibility pattern, the N210A substitution in VIM-2 had no substantial effect on the catalytic efficiencies against carbapenems, namely, imipenem and meropenem. Moreover, in the presence of zinc, VIM-2 and VIM-2_N210A exhibited nearly similar catalytic efficiencies against nitrocefin as observed in nitrocefin hydrolysis assay. The catalytic efficiency of VIM-2_E146A was slightly increased as compared to VIM-2. Hence, the *in vitro* catalytic efficiencies of VIM-2 and its mutants substantiated the antibiotic susceptibility pattern. Further, thermal stability in terms of melting temperature was determined for the purified proteins.

**Table 2.** Kinetic parameters of VIM-2 and its mutants for beta-lactam antibiotics.

| Antibiotics | Kinetic parameters | VIM-2 | Mutated VIM-2 variants |  |  |
| --- | --- | --- | --- | --- | --- |
|  |  |  | E146A | D182A | N210A |
| Ampicillin | $K_m$ ($\mu\text{M}$) | 76.5±5.52 | 69.3±6.07 | 117±18.5 | 94.1±5.78 |
| | $k_{\text{cat}}$ ($\text{s}^{-1}$) | 239±4.70 | 246±20.3 | 7.55±0.790 | 242±3.90 |
| | $k_{\text{cat}}/K_m$ ($\text{s}^{-1} \mu\text{M}^{-1}$) | 3.12±0.281 | 3.55±0.628 | 0.0744±0.0134 | 2.57±0.192 |
| Ticarcillin | $K_m$ ($\mu\text{M}$) | 188±35 | 85.4±7.25 | 134±4.98 | 206±28.6 |
| | $k_{\text{cat}}$ ($\text{s}^{-1}$) | 400±56.8 | 380±28.7 | 2.21±0.0137 | 207±34.8 |
| | $k_{\text{cat}}/K_m$ ($\text{s}^{-1} \mu\text{M}^{-1}$) | 2.13±0.109 | 4.44±0.702 | 0.0168±0.00132 | 1.01±0.0836 |
| Cephalothin | $K_m$ ($\mu\text{M}$) | 42.0±5.14 | 51.3±4.58 | 24.5±4.88 | 63.9±5.92 |
| | $k_{\text{cat}}$ ($\text{s}^{-1}$) | 99.0±9.78 | 161±15.1 | 2.85±0.167 | 49.1±2.22 |
| | $k_{\text{cat}}/K_m$ ($\text{s}^{-1} \mu\text{M}^{-1}$) | 2.36±0.0791 | 3.13±0.567 | 0.229±0.0614 | 0.791±0.103 |
| Cefotaxime | $K_m$ ($\mu\text{M}$) | 35.1±5.32 | 28.7±1.34 | 89.3±8.25 | 32.7±5.10 |
| | $k_{\text{cat}}$ ($\text{s}^{-1}$) | 81.0±7.26 | 73.7±11.9 | 6.41±0.326 | 15.5±0.172 |
| | $k_{\text{cat}}/K_m$ ($\text{s}^{-1} \mu\text{M}^{-1}$) | 2.32±0.142 | 2.56±0.543 | 0.0717±0.0111 | 0.473±0.0728 |
| Imipenem | $K_m$ ($\mu\text{M}$) | 20.9±2.07 | 18.5±3.22 | 18.6±3.62 | 23.0±2.82 |
| | $k_{\text{cat}}$ ($\text{s}^{-1}$) | 30.3±1.22 | 29.3±1.94 | 0.464±0.0313 | 33.0±1.19 |
| | $k_{\text{cat}}/K_m$ ($\text{s}^{-1} \mu\text{M}^{-1}$) | 1.46±0.106 | 1.58±0.357 | 0.0249±0.00559 | 1.44±0.221 |
| Meropenem | $K_m$ ($\mu\text{M}$) | 19.9±3.18 | 21.4±1.83 | 21.0±1.27 | 18.3±1.38 |
| | $k_{\text{cat}}$ ($\text{s}^{-1}$) | 17.9±1.96 | 18.7±0.657 | 0.297±0.00357 | 12.8±0.185 |
| | $k_{\text{cat}}/K_m$ ($\text{s}^{-1} \mu\text{M}^{-1}$) | 0.975±0.178 | 0.865±0.0495 | 0.0148±0.00998 | 0.689±0.0378 |
| Nitrocefin | $K_m$ ($\mu\text{M}$) | 63.3±11.2 | 47.5±6.02 | 17.18±0.88 | 61.8±6.02 |
| | $k_{\text{cat}}$ ($\text{s}^{-1}$) | 345±29.9 | 297±4.23 | 30.8±0.0376 | 297±3.59 |
| | $k_{\text{cat}}/K_m$ ($\text{s}^{-1} \mu\text{M}^{-1}$) | 5.45±1.43 | 6.23±0.515 | 1.79±0.107 | 4.79±0.516 |

### The mutations prominently affected the thermal stability of VIM-2

The stability of metallo β-lactamases is reportedly affected by the residues near the active-site (Bahr, *et al*. 2021, Kumar, *et al*. 2017). Metallo β-lactamases are most robust enzymes and their thermal stability has also been correlated with the enzyme activity. Therefore, we investigated the effect of E146A, D182A and N210A individual mutations on the thermal stability of VIM-2 (Fig. 3). All the proteins exhibited a small peak at around 30.5 °C which might be related to the dimerization of proteins (Fig. S2). VIM-2 exhibited unfolding that started from around 50 °C with a deep and narrower curve and the melting temperature was ∼58 °C. Though, the small peak at around 30.5 °C was evident for VIM-2_D182A, eventually the protein exhibited instability. VIM-2_N210A exhibited a reduction in the melting temperature by about 5.5 °C to ∼52.5 °C while the melting temperature of VIM-2_E146A was nearly equal to the wild type (∼60.5 °C). To correlate the effect of E146A, D182A and N210A substitutions on the melting temperatures and stability of VIM-2 with secondary structure stability of the proteins, the secondary structures were determined using circular dichroism spectroscopy (Fig. 3). The overall α-helicity of purified VIM-2 in presence of 50 µM zinc was observed to be 20.4%. However, in case of D182A substituted VIM-2, the α-helical structures were reduced to 12.1%. For N210A substitution, the α-helicity was reduced by ∼ 5% to about 15.4%, while VIM-2_E146A exhibited nearly 16.5% α-helicity. Therefore, it can be concluded D182A substitution leads to the instability of the while the N210A substitution affects the overall stability; though do not significantly affect the secondary structure.

**Fig. 3.**
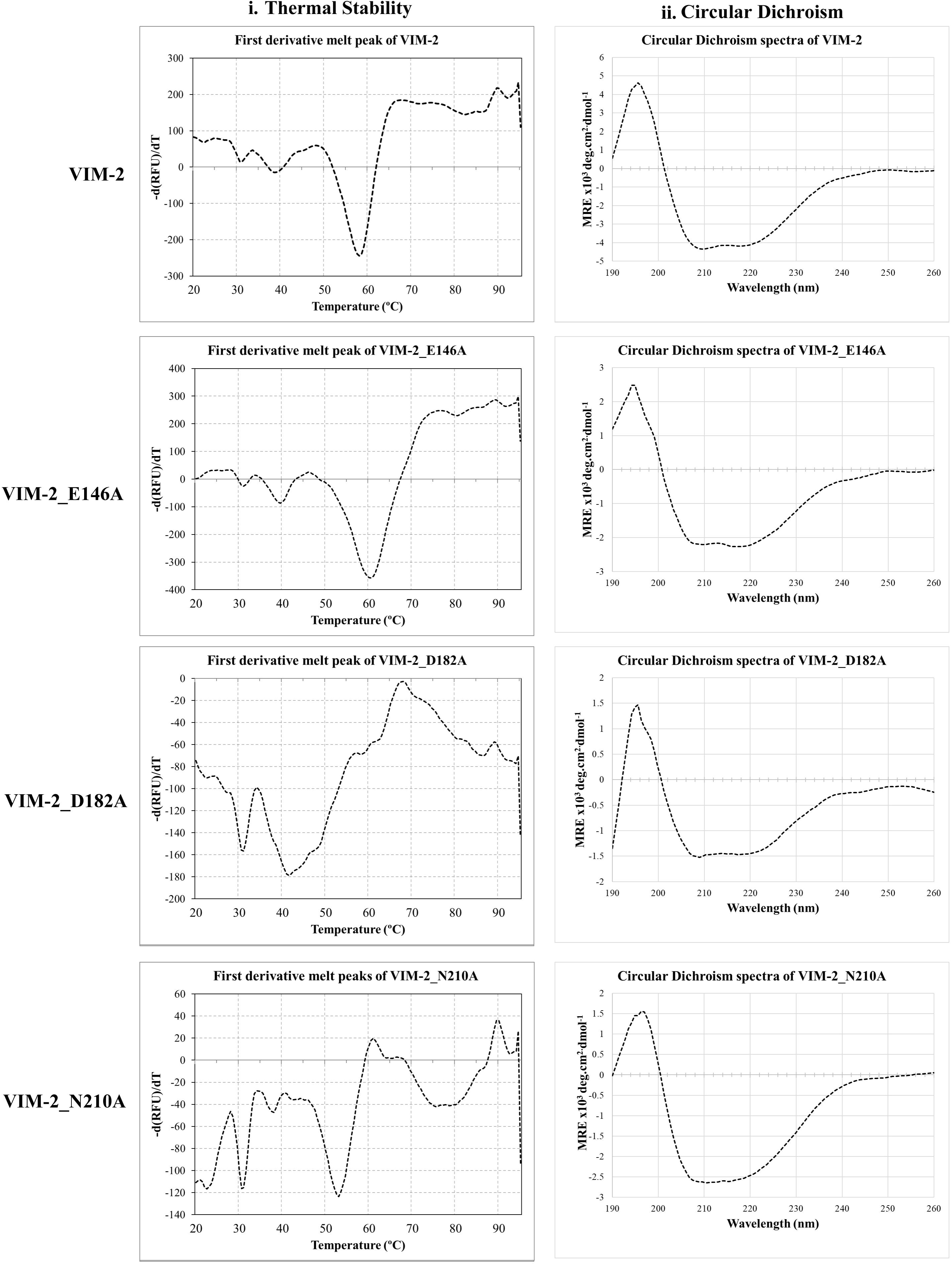
Assessment of thermal stability and secondary structures of VIM-2 and its mutants using differential scanning fluorimetry and circular dichroism.

## Discussion

The non-active-site residues play a significant role in maintaining the activity and stability of metallo β-lactamases (Borgianni, *et al*. 2010, Kumar, *et al*. 2017, Sun, *et al*. 2018). Interestingly, these residues, though conserved in some enzymes, exhibit variations in their functions (Bahr, *et al*. 2021) and can be involved in binding of inhibitors (Shin, et al. 2017). Therefore, in the present work we attempted to identify the role of conserved amino acid residues in VIM-2 that are reportedly involved in the activity of other metallo β-lactamases. As discussed previously, based on the literature survey the residues E146, D182, S207, N210 and D213 were substituted with alanine.

Out of these five residues, the expression of D182A and N210A substituted VIM-2, individually, in *E. coli* host, exhibited increased susceptibility to β-lactams in comparison to the wild-type VIM-2. The alanine substitution at residue D182 (BBL D199) drastically impacted the activity of VIM-2. This residue is conserved in nearly all the subclass B1 metallo β-lactamases and forms polar contacts with S180, T139 and main chain of Zn1 bound of H114 in VIM-2 (Fig. S3). D182 residue is a secondary residue for stabilizing the Zn1 in NDMs. Substitution of aspartate at this position with glutamate has resulted in about 4-fold reduction in the activity of NDM-1 (Sun, *et al*. 2018). Similarly, a reduction in the catalytic activity of NDM-1 with aspartate to alanine substitution at this position is also reported (Ali, *et al*. 2021). We observed a drastic reduction in both *in vivo* and *in vitro* activity of VIM-2 with D182A substitution. Also, the addition of zinc to purified VIM-2_D182A exhibited an improvement in the activity of VIM-2_D182A. The zinc ions are required for proper folding of metallo β-lactamases and the scarcity of zinc is correlated with the periplasmic instability of these enzymes (López, *et al*. 2022). The absence of aspartate at position 182 in VIM-2 leads to zinc deficiency which might be leading to periplasmic instability of the enzyme, thereby affecting its catalytic activity. The instability of this mutant was also evident by the thermal stability and the changes in the secondary structures. The residue N210 (BBL N233) showed substrate dependent variations in the activity of metallo β-lactamases, though conserved in most of the subclass B1 metallo β-lactamases. N210 residue bridges with Zn2 via a water molecule in VIM-2. The substitution of this asparagine residue with alanine in NDM-1 resulted in an attenuation of MICs for the ampicillin, carbapenems, and cephalosporins tested. The catalytic efficiency of this mutant is nearly similar to NDM-1 for ampicillin and lower catalytic efficiency was obtained for cefepime, imipenem, and ertapenem (Chiou, *et al*. 2014).

In IMP-1 substitution of this asparagine residue with alanine leads to an increased catalytic efficiency for the tested β-lactams (Materon and Palzkill 2001). However, another study on N233A substitution in IMP-1 has shown to decrease the catalytic efficiency of the protein against all tested β-lactams (Brown, *et al*. 2011).

Similar to these studies, the N210A (BBL N233A) substitution in VIM-2 has reduced the hydrolytic activity against tested penicillins and cephalosporins, while exhibited similar kind of activity against the tested carbapenems. Interestingly, the catalytic efficiency against cefotaxime was highly reduced. The reduced stability and the changes in the secondary structures can be attributed to the overall changes of polarity due to absence of polar asparagine residue in VIM-2_N210A. Although E146A substitution in VIM-2 imparted the activity and the stability comparable to VIM-2, further assessment of the mechanism of this residue is required for validating its role in the activity of this protein, since similar residue E152 is involved in the catalytic mechanism of NDMs (Kumar, *et al*. 2020). Overall, it can be inferred that the residues N210 and D182 individually play a critical role in the activity of VIM-2. All the observations are in line to our previous reports on the involvement of non-active site residues on the enzymatic activities of various β-lactamases. Therefore, it is important to look at these soft spots of various group of β-lactamases, which will help us design peptides or inhibitors that might inhibit the activity of these dreadful enzymes, and eventually, reduce the burden of β-lactam antibiotic resistance.

## Summary and Conclusion

Verona integron metallo β-lactamase (VIM-2) is one of the most widespread class B β - lactamase responsible for β-lactam resistance. Although active-site residues help in metal binding, the residues nearing the active-site possess functional importance and are crucial for comprehending the mechanism of its action. The expression of N210A substituted VIM-2 in the host displayed enhanced susceptibility towards tested penicillins and cephalosporins. D182A substitution had detrimental effect on VIM-2 activity. The host expressing VIM-2_N210A was susceptible to cefotaxime in the zinc depleting condition. Both the mutants displayed reduced thermal stability as compared to VIM-2. The secondary structures were altered for the mutants as compared to VIM-2. *In vitro* catalytic efficiency of VIM-2_N210A was reduced against penicillins and cephalosporins by >4 fold in the presence of zinc while D182A substitution in VIM-2 significantly reduced its activity. In a nutshell, N210A affects the substrate specificity and stability while D182A affect the overall stability of VIM-2.

## Acknowledgements

Authors are thankful to Central Research Facility, IIT Kharagpur for providing Circular Dichroism facility.

## Author’s contribution statement

DJ was involved in designing, experimentations, analysis and writing of the initial draft of the manuscript; TA was involved in experimentation and analysis; JV was involved in designing and experimentations; DC was involved in data analysis and edit of the manuscript and ASG was involved in conceptualization, designing, result analysis, writing and editing of the final draft of the manuscript.

