## Supplementary material for "Conserved ancillary residues situated proximally to the VIM-2 active site affect its metallo β-lactamase activity": Table S1; Table S2; Figure S1; Figure S2 and Figure S3

**Table S1.** Sequences of primers used for cloning and site directed mutagenesis

| Primer name | Primer sequence (5'-3' | Primer details |
|---|---|---|
| FP_VIM-2 | CTCTCTGAGCTCAGGAGGATATATATATGTTCAAACTTTTGAG | Cloning |
| RP_VIM-2 | CTCTCTAAGCTTCTACTCAACGACTGAGCG | |
| FP_sVIM-2 | CTCTCTCATATGTCCGTAGATTCTAGCGG | |
| FP_VIM-2_E146A | CGCCGGCTAGCCGAGGTAGCGGGGAACGAGATTCCCACG | Mutation |
| RP_VIM-2_E146A | CGTGGGAATCTCGTTCCCCGCTACCTCGGCTAGCCGGCG | |
| FP_VIM-2_D182A | GCTGCGCATTCGACCGCCAACTTAGTTGTGTAC | |
| RP_VIM-2_D182A | GTACACAACTAAGTTGGCGGTCGAATGCGCAGC | |
| FP_VIM-2_D213A | GCGGGGAACGTGGCCGCGGCCGATCTGGCTGAATGG | |
| RP_VIM-2_D213A | CCATTCAGCCAGATCGGCCGCGGCCACGTTCCCCGC | |
| FP_VIM-2_S207A | GAGTTGTCACGCACGGCTGCGGGGAACGTGGCC | |
| RP_VIM-2_S207A | GGCCACGTTCCCCGCAGCCGTGCGTGACAACTC | |
| FP_VIM-2_N210A | CGCACGTCTGCGGGGGCCGTGGCCGATGCCGATC | |
| RP_VIM-2_N210A | GATCGGCATCGGCCACGGCCCCCGCAGACGTGCG | |

**Table S2.** Parameters used for determination of catalytic efficiencies of the purified proteins

| Antibiotic | Concentration range (μM) | Molar extinction coefficient $\Delta\varepsilon$ ($M^{-1}cm^{-1}$) | Wavelength (nm) | Enzyme (nM) |
|---|---|---|---|---|
| **Penicillins** | | | | |
| Ampicillin | 250-1000 | -835 | 235 | 10-100 |
| Ticarcillin | 50-250 | -835 | 235 | 10-100 |
| **Cephalosporins** | | | | |
| Cefotaxime | 20-100 | -7500 | 260 | 10-100 |
| Cephalothin | 20-100 | -6500 | 260 | 10-100 |
| Nitrocefin | 20-100 | 15000 | 482 | 10-100 |
| **Carbapenems** | | | | |
| Imipenem | 10-50 | -9000 | 300 | 10-100 |
| Meropenem | 10-50 | -6500 | 300 | 10-100 |

**Fig. S1**

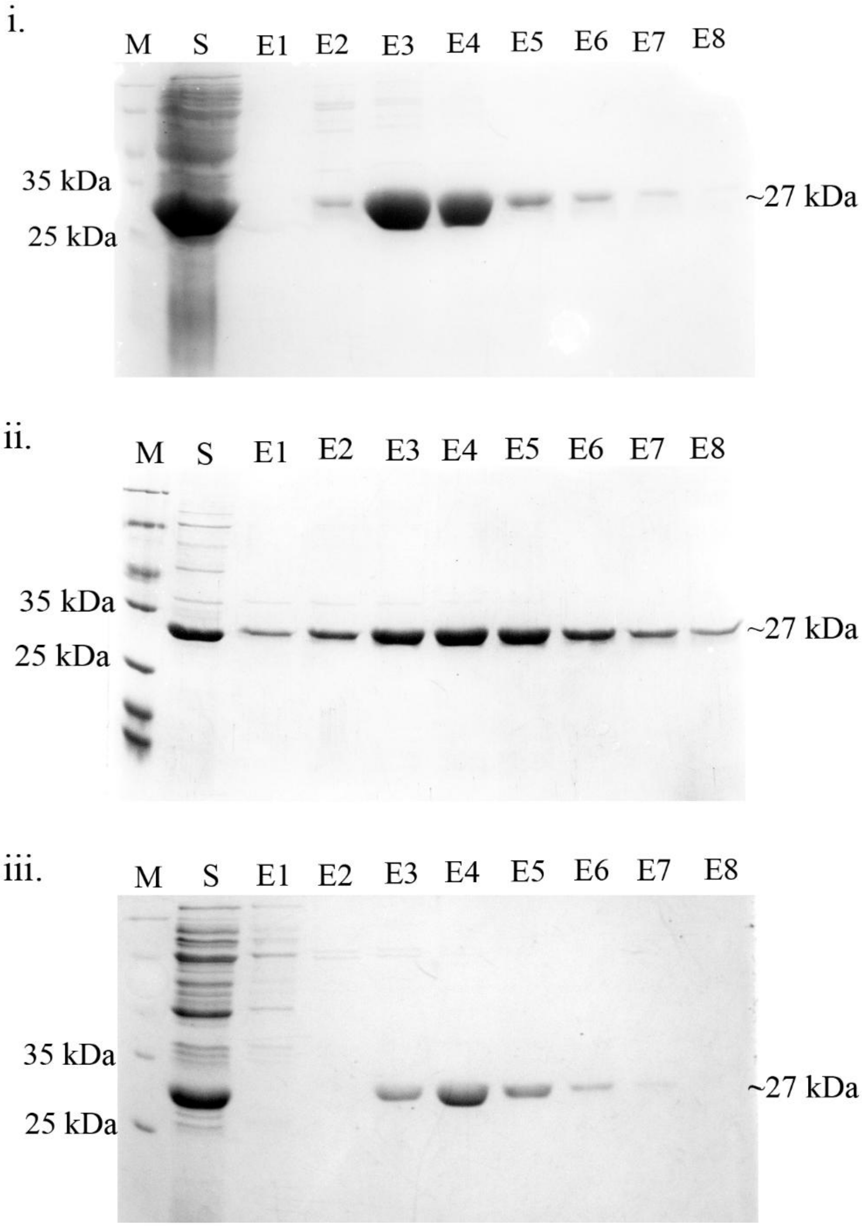


**Figure S1**. SDS-PAGE image of purification of VIM-2 and its mutants showing i. VIM-2 ii. VIM-2_D182A iii. VIM-2_N210A, where M stands for protein marker, S stands for the supernatant of *E. coli* BL21(DE3) cell lysates expressing proteins, E stands for the different elutes of purified protein.

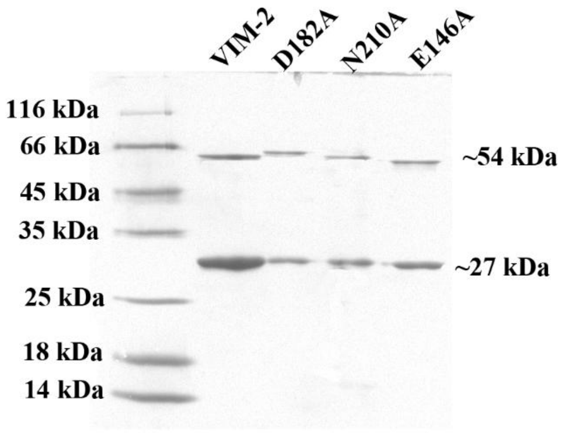


**Figure S2.** Native-PAGE image (Polyacrylamide Gel Electrophoresis without Sodium Dodecyl Sulphate) of VIM-2 and its mutants showing two bands at size 27 kDa and 54 kDa. The semi Native-PAGE was performed using 12% Tis-Glycine separating gel and 5% stacking gel devoid of SDS, with lower concentration of SDS (0.25%) in the running buffer. Sample loading dye did not contain SDS and reducing agent and samples were prepared without heating. The unstained protein marker used was also prepared without heating.

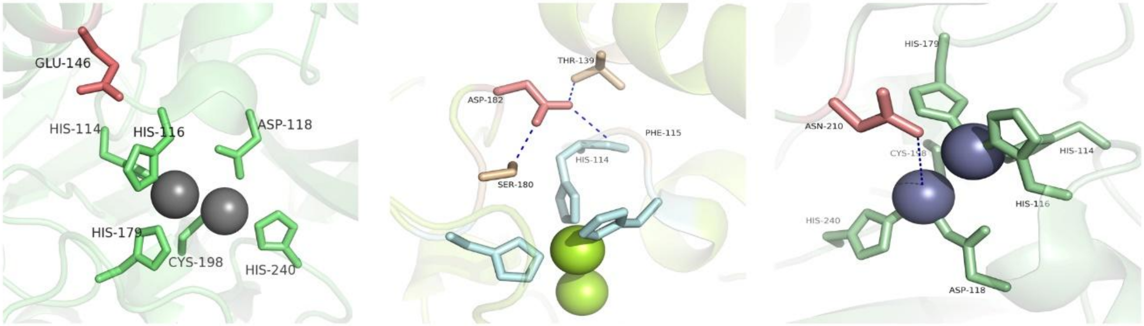


**E146** **D182** **N210**

**Figure S3.** The orientation of residues E146, D182 and N210 in deep salmon color with respect to active sites (PDB ID 4bz3).
